# Characterising the vegetation-rainfall relationship in the Northeast Himalaya, India

**DOI:** 10.1101/2021.10.19.464965

**Authors:** Bidyut Sarania, Vishwesha Guttal, Krishnapriya Tamma

**Author notes:** Centre for Ecological Sciences, Indian Institute of Science, Bengaluru.

## Abstract

Ecosystems are complex systems and are characterised by positive and negative feedbacks between the abiotic and biotic components. The response of an ecosystem to its environment can be determined by examining state diagrams, which are plots of the state variable as a function of the environmental driver. For instance, tree cover as a function of rainfall is widely used to characterise vegetation patterns. Previous studies have shown that tree cover shows bimodal distributions for intermediate rainfall regimes in Africa and South America. In this study, we construct a vegetation state diagram – by plotting vegetation cover as a function of mean annual rainfall – for Northeast India, which is part of the Eastern Himalaya and the Indo-Burma biodiversity hotspot. We use remotely sensed satellite data of Enhanced Vegetation Index (EVI) as a proxy for vegetation cover. We obtain Mean Annual Precipitation (MAP) from the CHIRPS data (Climate Hazards Group InfraRed Precipitation with Station data). We find that EVI increases monotonically as a function of MAP in the range 1000-2000 mm, after which it plateaus. The 1000 to 2000 mm MAP corresponds to the vegetation transitional zone (1200-3700 m), whereas >2000 MAP region covers the greater extent of the tropical forest (*<*1200 m) of NEI. In other words, we find no evidence for bimodality in tree cover or vegetation states at coarser scales in North Eastern India. Our characterisation of the state diagram for vegetation in northeast India is important to understand response to ongoing change in rainfall patterns.

## Introduction

An ecosystem is a complex dynamical system comprising of various interacting, strongly interdependent biotic and abiotic components, with both positive and negative feedbacks between them (Scheffer et al. 2001; Scheffer and Carpenter 2003). The existence of features such as feedback loops and stochasticity - both environmental and demographic – makes it difficult to predict responses of ecosystems to changing environmental driver variables.

To understand how ecosystems respond to changing driver, we can plot of the state variable of ecosystem as a function of the environmental driver, called a state diagram (Hirota et al. 2011; Majumder et al. 2019). Vegetation cover, coral density, population density, plankton diversity are examples of macroscopic state variables that can be used to characterise various ecological systems, and to investigate how they respond to environmental drivers (Staver et al. 2011; Tanzil et al. 2013; Bonecker et al. 2013). As a specific and widely studied example of state diagram in the context of biomes, understanding relationship between vegetation and mean annual precipitation have helped ecologists characterise the response of these systems to spatial and temporal variation in rainfall (Sankaran et al. 2005; Staal et al. 2016; Majumder et al. 2019).

In general, state diagrams can be of different forms showing linear, non-linear, and bistable relationships between ecosystem state and the environmental drivers. A system that shows a linear relationship will likely show a gradual (more predictable) response to changing environmental driver (Feher et al. 2017). On the other hand, in bistable ecological systems the ecosystem state can transition from one state to an alternative state when the driver crosses a threshold value (Scheffer and Carpenter 2003) or when stochasticity pushes the system beyond the basin of attraction of the current state (Guttal and Jayaprakash 2007). Such transitions, also called regime shifts, tipping points or critical transitions, can dramatically alter the structure and functioning of these ecosystems. It is now well established that mean annual precipitation is an important driver of tree cover, globally (Hirota et al. 2011; Staver et al. 2011; Staal et al. 2016; Majumder et al. 2019). The nature of the relationship between vegetation cover and mean annual precipitation is complex, and is not similar across continents (Xu et al. 2018). Previous studies have shown that tree cover in Africa (and parts of Australia and South America) show multiple stable states, with potential for abrupt transitions and hysteresis behaviour between the alternative states (Staver et al. 2011). Such characterisations, via state diagrams, are important to understand the response of vegetation cover to mean annual precipitation (MAP) as climate change and other human interventions threaten the forest and savanna ecosystems (Staal et al. 2020).

Despite the presence of forests (tropical, subtropical), grasslands and mesic savannas, no such characterization has been carried out for Indian ecosystems (Ratnam et al. 2016). Mean annual precipitation in India shows strong spatial gradients, which in turn influences the diversity of vegetation types (Roy et al. 2015). Tropical and sub-tropical forests (evergreen, semi-evergreen, moist deciduous) are typically found in areas of high rainfall (>1500 mm), whereas dry deciduous, thorny and scrub forest are typically found in areas of low rainfall (*<*1000 mm) (Reddy et al. 2015). Interestingly, forest-grassland complexes such as tropical montane forests-grasslands of Western Ghats (Shola forest) also occur. Here, forests-grasslands exist as alternative states, with frost and freezing temperatures maintaining the two stable states (Joshi et al. 2020). Apart from this, other vegetation mosaics have also been reported, including woodland-grassland of western India (Kumar et al. 2015), and forest-grasslands / shrublands of the Himalayas (Singh and Singh 1987; Dvorsky’ et al. 2011; Rawat 2017; Yadava 1990). In this study, we focus on vegetation patterns in northeast India (henceforth NEI) which is a part of two biodiversity hotspots - the Indo-Malayan and the eastern Himalaya (Myers et al. 2000). The eastern Himalayas along with the hills of NEI (including the Garo-Khasi-Jaintia hills, Arakan hills) also show tremendous diversity in vegetation, and rainfall. This region encompasses a wide elevational range from 20 m to >6000 m above mean sea level, resulting in a large climate gradient that ranges from tropical, sub-tropical, temperate to alpine (Dikshit and Dikshit 2014). The NEI region receives the highest rainfall during the south-west monsoon of the Indian sub-continent, but the amount of rainfall at local scales is highly variable due to the rain shadow effects (Dikshit and Dikshit 2014; Prokop and Walanus 2015). The steep elevational gradient, along with the heterogeneity in rainfall across NEI, results in various vegetation types, including tropical, subtropical and temperate forests, grasslands (both cold-arid, and flooded) and savanna (Champion and Seth 1968; Acharya et al. 2011; Yadava 1990; Saikia et al. 2017). The ecosystems of NEI are amongst the most vulnerable to climate change, with many parts already showing changes in climate patterns, for instance, trends of rainfall (Oza et al. 2014; Preethi et al. 2017). This also has enormous impacts on the local communities that live in close association with these landscapes. Characterising the state diagram for vegetation in this region is therefore essential for long-term monitoring and conservation of these systems.

In this paper, we construct the state diagram of forest ecosystems of Northeastern India to characterise the relationship between vegetation cover and mean annual precipitation. Since the region also shows a large elevational gradient, we also examine the relationship between vegetation cover and elevation to determine its role in driving patterns.

## Methods

### Study Site

Northeast India (NEI) consists of eight states (Sikkim, Assam, Meghalaya, Arunachal Pradesh, Nagaland, Mizoram, Tripura, and Manipur), and comprises of 7.9 % of India’s total land mass (Fig. 1). The states of Sikkim, Arunachal Pradesh, and the northernmost part of Assam lies in the eastern Himalaya biogeographic realm, while the southernmost parts of Assam, and state of Meghalaya, Mizoram, Tripura, Manipur, and Nagaland lies in Indo-Malayan biogeographic realm.

**Figure 1:**
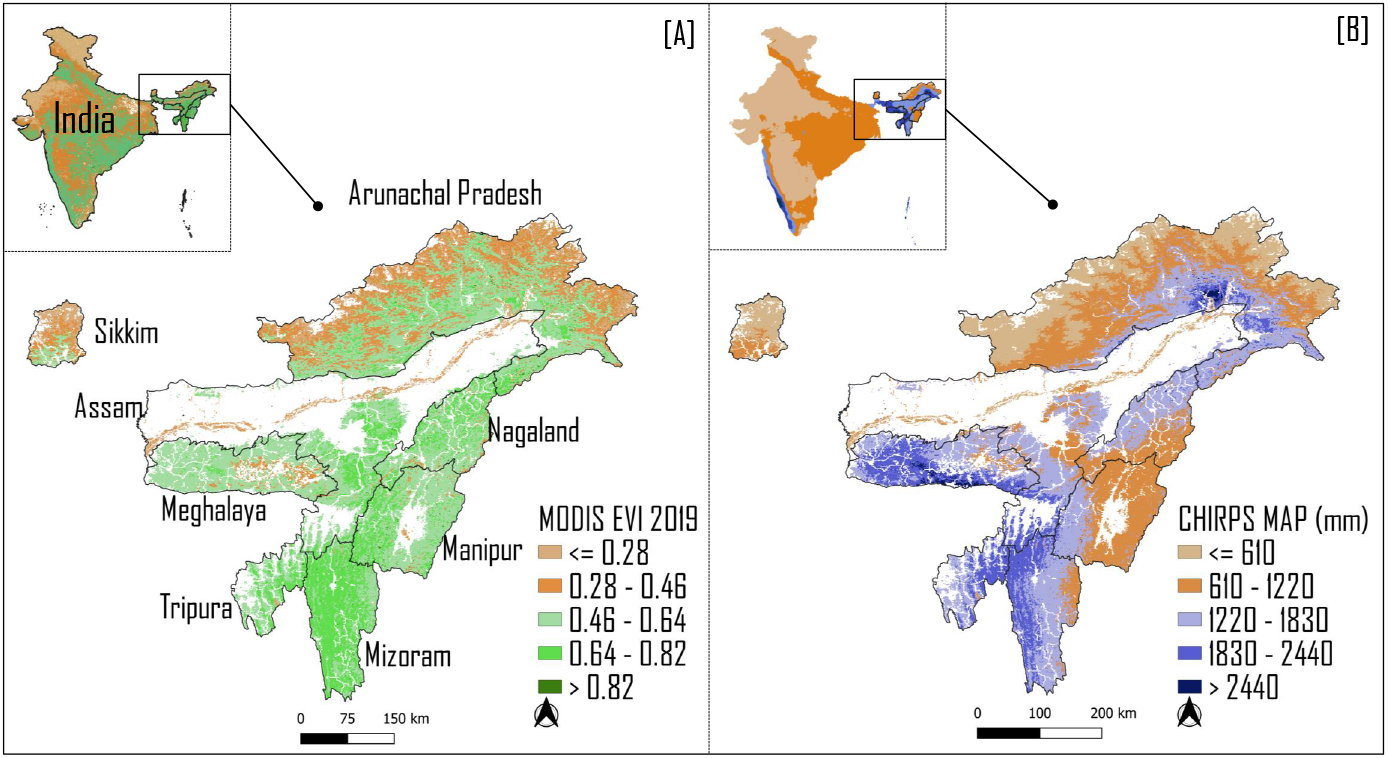
Map of Northeast India showing spatial patterns of [A] Enhanced Vegetation Index, and [B] Mean Annual Precipitation. From both these images, pixels with high human footprint have been removed (see methods for details).

### Vegetation cover data

To characterise the state of the ecosystem, we considered vegetation cover as a state variable and mean annual precipitation (MAP) as the driver (Hirota et al. 2011; Majumder et al. 2019). We used the Enhanced Vegetation Index (EVI), obtained from the Moderate Resolution Imaging Spectroradiometer (MODIS), as a proxy for vegetation cover. We obtained EVI at 1 km resolution obtained from MODIS (dataset: MCD43A4), using the Google Earth Engine platform (Huete et al. 2002; Gorelick et al. 2017). We constructed a composite based on images obtained in the post-monsoon season (1st October, 2019-30th November, 2019). This period was chosen as the vegetation cover reflects the highest possible in the growing season, while minimising cloud cover. EVI is an atmospherically corrected index, is less sensitive to soil background, and is sensitive to canopy variation (Huete et al. 2010). Thus, EVI is well suited for tropical regions with high biomass compared to other vegetation indices such as normalised difference vegetation index (NDVI) and vegetation continuous fields (VCF)(Huete et al. 2002).

### Rainfall and Elevation Data

We obtained rainfall data for the period spanning 1989-2019 from Climate Hazards Group InfraRed Precipitation dataset (CHIRPS), at a spatial resolution of 0.05 arc degrees (Funk et al. 2015). We calculated mean annual precipitation (MAP), aggregated it to a final resolution of 1 km prior to analyses. We obtained elevation data at a spatial resolution of 30 m from the Shuttle Radar Topography Mission (SRTM) data (Farr et al. 2007). The data was rescaled to a 1 km final resolution prior to analyses. We used Global terrestrial Human Footprint maps (HFI) and Glob-Cover maps as a masks to eliminate pixels with high human footprint, especially associated with built-up and agricultural areas (Majumder et al. 2019; Bontemps et al. 2011; Venter et al. 2016). The HFI value of NEI ranges from 0-43, and typically most pixels associated with natural vegetation lie within the range of > 0 to ≤ 8 (See Supplementary 1). Thus, we removed pixels with HFI values > 8 for EVI data. We also analysed the data without removing the human footprint (see Supplementary 2). All datasets were processed and analyzed via the Google Earth Engine platform.

### State Diagram

State diagram is a plot of the vegetation cover (EVI) as a function of the mean annual rainfall (Hirota et al. 2011; Majumder et al. 2019). Following the method of Mazumdar et al (2019), we divided the mean annual rainfall into 100 mm bins. For each bin, we obtained the corresponding EVI data, and constructed a smoothed frequency distribution of the same using the density function in R (Majumder et al. 2019). From this, we identified the modes (local maxima) of distribution. To consider a local maxima as a mode, we followed the protocol of Majumder et al. (2019). A local maxima is considered as a distinct mode if it satisfies two conditions: a) the ratio between two local maxima is greater than 0.25, and b) the distance the two local maxima is greater than 0.1 EVI units. If the ratio 0.25 but the distance between the mode is ≤ 0.1, then the weighted average of modes is taken. This approach has been adopted to avoid small peaks that appears due to unknown stochastic reasons (Majumder et al. 2019). The sorted EVI modes are then plotted as a function of annual mean precipitation. To determine the influence of elevation on vegetation patterns, we also constructed a state diagram of EVI with elevation as the driver. Following the same protocol (described above), we divided elevation into 100 m bins, and obtained the modes of EVI for each bin.

## Results

We display the state diagram - which is a plot of EVI as function of mean annual precipitation (MAP) - for the NEI in Fig 2. From our plot, we observed that EVI increases linearly with MAP up to 2000 mm, beyond which there is no further increase in EVI. We did not find evidence for bimodality in EVI for any rainfall regime, which suggests that the system does not show bistability (or the presence of alternate stable states).

**Figure 2:**
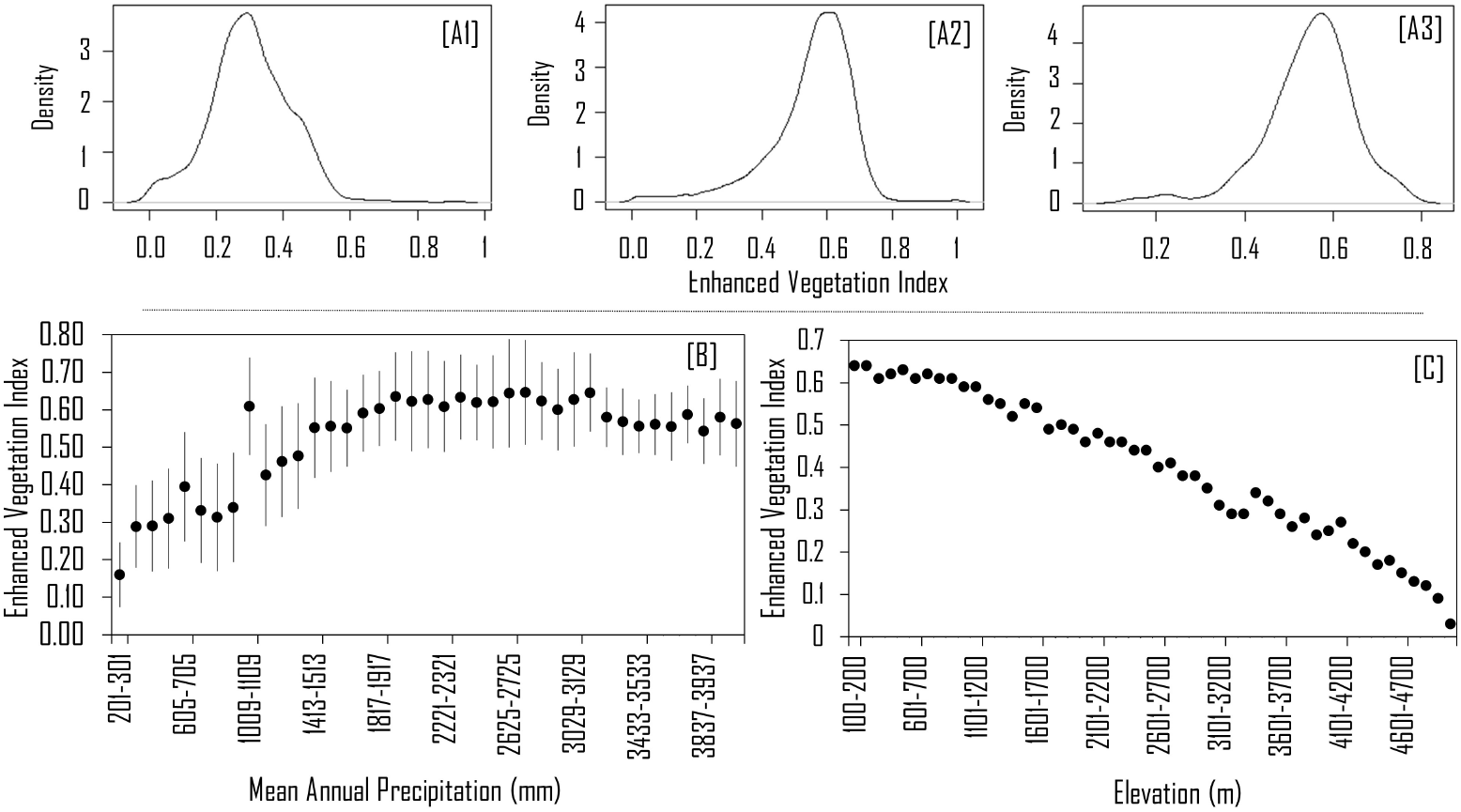
Frequency distribution of EVI (smoothed showing density function in R) for different rainfall bins corresponding to [A1] 403-503 mm, [A2] 1009-1109 mm, [A3] 4039-4139 mm. The observed distributions are all unimodal. [B] State diagram, where mode (±SD) of EVI increased gradually with MAP. [C] Mode of EVI decreased monotonically with increase in elevation. The rainfall bins are 100mm bins, while the elevation bins are 100m bins.

To determine the influence of elevation in driving the initial increase of EVI, we constructed the state diagram for EVI as a function of elevation. We found that EVI decreased monotonously with increase in elevation (Fig. 2 C). Additionally, our analysis showed that temperature and mean annual precipitation also decreased monotonously with elevation (Supplementary 4). To investigate this relationship further, we examined the relationship of three large bands of precipitation (<1000 mm, 1000-2000 mm and >2000 mm) with elevation and vegetation cover (Fig. 3), as these bands had distinct frequency distributions of EVI in the state diagram. Regions with MAP ≤ 1000 mm showed an elevation profile > 2500 m (Fig. 3 A and B), while regions with MAP > 2000 mm typically show elevation profile < 1000 m (Fig. 3 D and A). In other words, high elevation regions typically showed lower MAP, while low elevation regions were associated with higher MAP. On the other hand, the regions with 1000-2000 mm MAP includes elevational gradient that spans between 1200-3700 m (Fig. 3 B and A).

**Figure 3:**
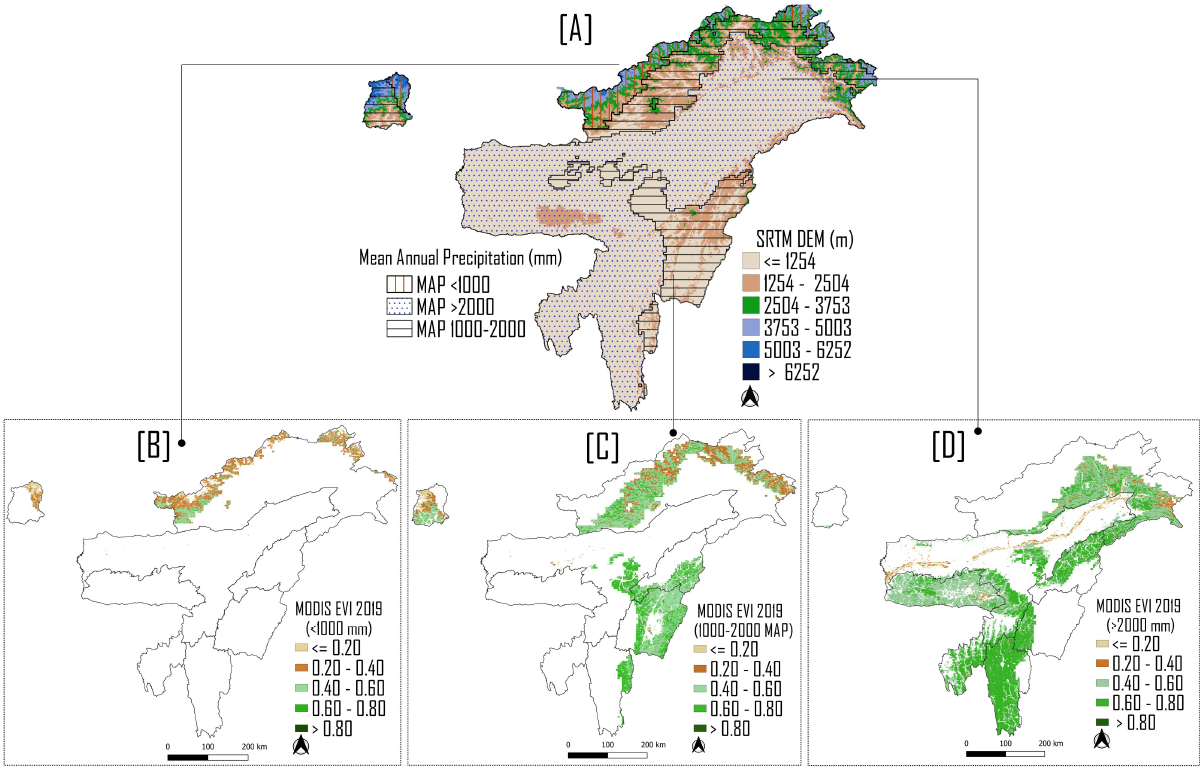
[A] Map of Elevation (SRTM DEM) and spatial distribution of EVI for [B] MAP of <1000 mm, [C] 1000-2000 mm, and [D] >2000 mm. Elevational gradients map shows that MAP of <1000 mm corresponds to elevation of >2500 m, 1000-2000 mm corresponds to >1200-3700 m elevation, and >2000 mm MAP corresponds to <1200 m elevation

## Discussion

In this study, we constructed the state diagram for vegetation cover, by characterising EVI as a function mean annual precipitation (MAP) for northeast India (NEI). The state diagram shows that the EVI gradually increases with MAP, and reaches an asymptote at values greater than 2000 mm. Throughout this range of rainfall values, we did not find evidence for bimodal distribution of vegetation cover, indicating absence of bistable forest states in NEI. We also found that EVI monotonically decreases as elevation increases; furthermore, higher elevations are associated with reduced MAP and lower temperatures, both factors possibly contributing to lower EVI. This is consistent with other studies showing a decrease in tree species richness as elevation increases (Acharya et al. 2011; Behera and Kushwaha 2006). Our study provides the first assessment of coarse scale relationship between vegetation cover and rainfall for northeastern India in the context of forest resilience and multiple stable states.

We note the contrast with the characteristics of coarse-scale vegetation in Africa, South America and some parts of Australia, which exhibit bimodal distribution of tree cover at intermediate mean annual rainfall regimes corresponding to 1000 mm to 2500 mm (Staver et al. 2011; Hirota et al. 2011; Staal et al. 2016; Majumder et al. 2019). Although, there is no definite pattern of biomodality observed for NEI, there is a monotonic increase in vegetation cover (EVI) at intermediate rainfall regimes (1000 - 2000 mm of MAP). This intermediate rainfall regime corresponds to a vegetation transitional zone (1200-3700 m) of NEI, where we find a variety of forests: subtropical broadleaved forest at >1200-2000 m, mixed coniferous forest or temperate forest at >2000-3000 m, and sub-alpine vegetation types at elevation of >2900 m (Saikia et al. 2017). As a result, the monotonic increase in EVI at intermediate MAP could be due to the presence of different vegetation types. On the other hand, at > 2000 mm MAP, typically tropical evergreen and semi-evergreen forests are dominant, which explains the plateau observed in the state diagram (Champion and Seth 1968).

Although, we did not find bimodality in NEI at the coarse scale (1 sq. km), we do not rule out that small patches of forest-grassland mosaics occur. Indeed, several small pockets of grassland and savanna are found in the hills and valleys of NEI (Yadava 1990; Choudhury et al. 2017), including Dzuku valley, Balpakram NP, Kaziranga NP and Manas NP, apart from the high elevation grasslands (Yadava 1990; Mistry and Stott 1993; Choudhury et al. 2017). The natural grassland and savanna ecosystems in NEI occupy a smaller extent compared to the forest ecosystems. In addition, the savanna and grasslands of the Brahmaputra valley have been altered extensively for agricultural activity. This has further reduced the extent of these ecosystems in the landscape. The presence of forest-grassland mosaics, albeit at very small spatial scales, suggests the presence of mechanisms that maintain such mosaics. We argue that fine scale analysis with high resolution data is necessary to understand the dynamics and stability of such mosaic landscapes.

Savannas cover a large area in Africa at intermediate rainfall levels, and the distribution of this bistable system is likely maintained by a complex interplay of fire, herbivores and dispersal (Sankaran et al. 2005; Murphy and Bowman 2012; Vanak et al. 2012). For instance, as the amount of grass biomass increases, fires become more intense, limiting forest growth, whereas decreases of grass biomass result in less fire that allows forest to expand. Besides that, herbivores play an important role in the fire-grass interaction, as the degree of grazing affects the load of grass biomass.Slash and burn cultivation (shifting cultivation) is the major reason for the forest fire in the NEI; however the effect of fire and herbivores on the forest’s stability remains unclear (Yadava 1990). Flood, on the other hand, is known to have a positive feedback in maintaining floodplain grassland ecosystem of NEI’s, such as Assam’s Kaziranga NP (Khatri et al. 2011). There is limited knowledge on the interplay of fire, grazing, and flood in maintaining forest-savanna ecosystems in NEI, which therefore needs further attention.

Vegetation in the region has responded to past changes in climate. Paleoecological investigations of the recent past – up to 1200 years before present – show that the climate of NEI has been relatively stable (Agrawal et al. 2015), possibly, allowing for stability in vegetation structure and composition. However, over longer time frames, such as that of the Holocene, the climate in NEI has likely fluctuated between cool, dry and warm, moist conditions (Bhattacharyya et al. 2007). Such past climate fluctuations have likely driven changes in vegetation. For instance, paleoecological analyses from Assam provide evidence for shifts between savanna and tropical mixed deciduous forests over the past 10,000 years, including instances of floods in the valley (Dixit and Bera 2013; Mehrotra et al. 2014). Changes in vegetation patterns in response to changes in precipitation have also been reported from other regions of Eastern Himalayas (Bhattacharyya et al. 2014), Garo hills of Meghalaya (Basumatary et al. 2014) and from the Arakan range of Nagaland (Misra et al. 2020). Therefore, a characterisation of relationship between vegetation cover and rainfall along with palaeological information is important for predicting future changes in vegetation pattern.

Indeed, ongoing climate change in the Himalayas is causing changes in precipitation patterns, seasonality, and temperature (Singh et al. 2016). The NEI region is already experiencing rainfall deficit and is expected to face drought as a result of future climate change (Choudhury et al. 2019). Vegetation response to these changes are likely to be complex and spatially heterogeneous. Although data on the response of vegetation to the ongoing climate change is sparse from the Eastern Himalayas, palaeological data, along with ongoing ecological studies demonstrate how ongoing and future climate changes in the region can lead to shifts in vegetation, biodiversity loss, and loss of ecosystem services in the region (Xu et al. 2009). Therefore, a further decrease of MAP (<1000 mm), an increase in incidence of forest fires, and deforestation can lead to shifts in vegetation (such as, forest to savanna) or change in the plant community structure. Such changes have indeed been observed in other parts tropical regions, for instance, in the savanna of Africa (Sankaran et al. 2005), and in Amazonian rain forests (Ziccardi et al. 2021). Therefore, further studies exploring the impact of fires, changing rainfall patterns, deforestation on the NEI forests are needed.

While our study has focused on northeast India, there is a need for such an analysis at the scale of the Indian subcontinent. The region has a diversity of ecosystems including savanna, scrub, and tropical forests. Such an investigation will provide us with more insights into the structuring of vegetation in the subcontinent.

In summary, our study provides a first coarse-scale characterisation of how vegetation changes with mean annual precipitation. Our work may aid better understanding of vulnerability of ecosystems in the Eastern Himalayas to undergo transitions into alternative states, in the context of large scale climate change as well as local anthropogenic drivers, especially in regions receiving 1000 to 2000 mm of mean annual precipitation. We argue that there is an urgent need for long-term monitoring of ecosystems at both coarse and fine scales in these regions. Furthermore, an extension of such a vegetation characterisation to other parts of India is also desirable.

## Acknowledgements

We acknowledge the Department of Biotechnology and DBT-IISc Partnership Program for support. We thank ArunKumar Mahendran and Nainika Konwar for the technical support.

## References

B. K. Acharya, B. Chettri, and L. Vijayan. Distribution pattern of trees along an elevation gradient of eastern himalaya, india. Acta Oecologica, 37(4):329–336, 2011.

S. Agrawal, P. Srivastava, N. Meena, S. K. Rai, R. Bhushan, D. Misra, A. Gupta, et al. Stable (δ 13 c and δ 15 n) isotopes and magnetic susceptibility record of late holocene climate change from a lake profile of the northeast himalaya. Journal of the Geological Society of India, 86(6): 696–705, 2015.

S. Basumatary, S. Bera, S. Sangma, and G. Marak. Modern pollen deposition in relation to vegetation and climate of balpakram valley, meghalaya, northeast india: Implications for indo-burma palaeoecological contexts. Quaternary International, 325:30–40, 2014.

M. D. Behera and S. P. S. Kushwaha. An analysis of altitudinal behavior of tree species in subansiri district, eastern himalaya. In Plant Conservation and Biodiversity, pages 277–291. Springer, 2006.

A. Bhattacharyya, J. Sharma, S. K. Shah, and V. Chaudhary. Climatic changes during the last 1800 yrs bp from paradise lake, sela pass, arunachal pradesh, northeast himalaya. Current Science, pages 983–987, 2007.

A. Bhattacharyya, N. Mehrotra, S. K. Shah, N. Basavaiah, V. Chaudhary, and I. B. Singh. Analysis of vegetation and climate change during late pleistocene from ziro valley, arunachal pradesh, eastern himalaya region. Quaternary Science Reviews, 101:111–123, 2014.

C. C. Bonecker, N. R. Simões, C. V. Minte-Vera, F. A. Lansac-Tôha, L. F. M. Velho, and Â. A. Agostinho. Temporal changes in zooplankton species diversity in response to environmental changes in an alluvial valley. Limnologica, 43(2):114–121, 2013.

S. Bontemps, P. Defourny, E. Van Bogaert, O. Arino, V. Kalogirou, and J. R. Perez. Globcover 2009-products description and validation report. URL: http://ionia1.esrin.esa.int/docs/GLOBCOVER2009_Validation_Report 2, 2, 2011.

H. G. Champion and S. K. Seth. A revised survey of the forest types of India. Manager of publications, 1968.

B. A. Choudhury, S. K. Saha, M. Konwar, K. Sujith, and A. Deshamukhya. Rapid drying of northeast india in the last three decades: Climate change or natural variability? Journal of Geophysical Research: Atmospheres, 124(1):227–237, 2019.

M. R. Choudhury, P. Deb, H. Singha, and B. Chakdar. Structure and composition of a protected dry savanna in the northern brahmaputra floodplains of assam, northeast india. Range Management and Agroforestry, 38(1):27–34, 2017.

K. Dikshit and J. K. Dikshit. Weather and climate of north-east india. In North-East India: land, people and economy, pages 149–173. Springer, 2014.

S. Dixit and S. Bera. Pollen-inferred vegetation vis-à-vis climate dynamics since late quaternary from western assam, northeast india: Signal of global climatic events. Quaternary International, 286:56–68, 2013.

M. Dvorsky’, J. Doležal, F. De Bello, J. Klimešovà, and L. Klimeš. Vegetation types of east ladakh: species and growth form composition along main environmental gradients. Applied Vegetation Science, 14(1):132–147, 2011.

T. G. Farr, P. A. Rosen, E. Caro, R. Crippen, R. Duren, S. Hensley, M. Kobrick, M. Paller, E. Rodriguez, L. Roth, et al. The shuttle radar topography mission. Reviews of geophysics, 45(2), 2007.

L. C. Feher, M. J. Osland, K. T. Griffith, J. B. Grace, R. J. Howard, C. L. Stagg, N. M. Enwright, K. W. Krauss, C. A. Gabler, R. H. Day, et al. Linear and nonlinear effects of temperature and precipitation on ecosystem properties in tidal saline wetlands. Ecosphere, 8(10):e01956, 2017.

C. Funk, P. Peterson, M. Landsfeld, D. Pedreros, J. Verdin, S. Shukla, G. Husak, J. Rowland, L. Harrison, A. Hoell, et al. The climate hazards infrared precipitation with stations—a new environmental record for monitoring extremes. Scientific data, 2(1):1–21, 2015.

N. Gorelick, M. Hancher, M. Dixon, S. Ilyushchenko, D. Thau, and R. Moore. Google earth engine: Planetary-scale geospatial analysis for everyone. Remote sensing of Environment, 202: 18–27, 2017.

V. Guttal and C. Jayaprakash. Impact of noise on bistable ecological systems. Ecological modelling, 201(3-4):420–428, 2007.

M. Hirota, M. Holmgren, E. H. Van Nes, and M. Scheffer. Global resilience of tropical forest and savanna to critical transitions. Science, 334(6053):232–235, 2011.

A. Huete, K. Didan, T. Miura, E. P. Rodriguez, X. Gao, and L. G. Ferreira. Overview of the radiometric and biophysical performance of the modis vegetation indices. Remote sensing of environment, 83(1-2):195–213, 2002.

A. Huete, K. Didan, W. van Leeuwen, T. Miura, and E. Glenn. Modis vegetation indices. In Land remote sensing and global environmental change, pages 579–602. Springer, 2010.

A. A. Joshi, J. Ratnam, and M. Sankaran. Frost maintains forests and grasslands as alternate states in a montane tropical forest–grassland mosaic; but alien tree invasion and warming can disrupt this balance. Journal of Ecology, 108(1):122–132, 2020.

P. Khatri, K. Barua, et al. Structural composition and productivity assessment of the grassland community of kazhiranga national park, assam. Indian Forester, 137(3):290, 2011.

V. Kumar, A. Mahato, and R. Patel. Ecology and management of banni grassland of kachchh gujarat. Ecology and management of Grassland Habitats in India, ENVIS Bulletin: wildlife and protected areas, Wildlife Institute of India, Dehradun, 17:42–53, 2015.

S. Majumder, K. Tamma, S. Ramaswamy, and V. Guttal. Inferring critical thresholds of ecosystem transitions from spatial data. Ecology, 100(7):e02722, 2019.

N. Mehrotra, S. K. Shah, and A. Bhattacharyya. Review of palaeoclimate records from northeast india based on pollen proxy data of late pleistocene–holocene. Quaternary International, 325: 41–54, 2014.

S. Misra, S. Bhattacharya, P. K. Mishra, K. G. Misra, S. Agrawal, and A. Anoop. Vegetational responses to monsoon variability during late holocene: Inferences based on carbon isotope and pollen record from the sedimentary sequence in dzukou valley, ne india. Catena, 194:104697, 2020.

J. Mistry and P. Stott. The savanna forests of manipur state, india: an historical overview. Global ecology and biogeography letters, pages 10–17, 1993.

B. P. Murphy and D. M. Bowman. What controls the distribution of tropical forest and savanna? Ecology letters, 15(7):748–758, 2012.

N. Myers, R. A. Mittermeier, C. G. Mittermeier, G. A. Da Fonseca, and J. Kent. Biodiversity hotspots for conservation priorities. Nature, 403(6772):853–858, 2000.

M. Oza, C. Kishtawal, et al. Spatial analysis of indian summer monsoon rainfall. J. Geomatics, 8 (1):41–47, 2014.

B. Preethi, M. Mujumdar, R. Kripalani, A. Prabhu, and R. Krishnan. Recent trends and teleconnections among south and east asian summer monsoons in a warming environment. Climate Dynamics, 48(7-8):2489–2505, 2017.

P. Prokop and A. Walanus. Variation in the orographic extreme rain events over the meghalaya hills in northeast india in the two halves of the twentieth century. Theoretical and Applied Climatology, 121(1):389–399, 2015.

J. Ratnam, K. W. Tomlinson, D. N. Rasquinha, and M. Sankaran. Savannahs of asia: antiquity, biogeography, and an uncertain future. Philosophical Transactions of the Royal Society B: Biological Sciences, 371(1703):20150305, 2016.

G. S. Rawat. 12 the himalayan vegetation along horizontal and vertical gradients. Bird Migration Across the Himalayas: Wetland Functioning amidst Mountains and Glaciers, page 189, 2017.

C. S. Reddy, C. S. Jha, P. Diwakar, and V. K. Dadhwal. Nationwide classification of forest types of india using remote sensing and gis. Environmental monitoring and assessment, 187(12):1–30, 2015.

P. S. Roy, M. D. Behera, M. Murthy, A. Roy, S. Singh, S. Kushwaha, C. Jha, S. Sudhakar, P. K. Joshi, C. S. Reddy, et al. New vegetation type map of india prepared using satellite remote sensing: Comparison with global vegetation maps and utilities. International Journal of Applied Earth Observation and Geoinformation, 39:142–159, 2015.

P. Saikia, J. Deka, S. Bharali, A. Kumar, O. Tripathi, L. Singha, S. Dayanandan, and M. Khan. Plant diversity patterns and conservation status of eastern himalayan forests in arunachal pradesh, northeast india. Forest Ecosystems, 4(1):1–12, 2017.

M. Sankaran, N. P. Hanan, R. J. Scholes, J. Ratnam, D. J. Augustine, B. S. Cade, J. Gignoux, S. I. Higgins, X. Le Roux, F. Ludwig, et al. Determinants of woody cover in african savannas. Nature, 438(7069):846–849, 2005.

M. Scheffer and S. R. Carpenter. Catastrophic regime shifts in ecosystems: linking theory to observation. Trends in ecology & evolution, 18(12):648–656, 2003.

M. Scheffer, S. Carpenter, J. A. Foley, C. Folke, and B. Walker. Catastrophic shifts in ecosystems. Nature, 413(6856):591–596, 2001.

J. Singh and S. Singh. Forest vegetation of the himalaya. The Botanical Review, 53(1):80–192, 1987.

S. Singh, R. Kumar, A. Bhardwaj, L. Sam, M. Shekhar, A. Singh, R. Kumar, and A. Gupta. Changing climate and glacio-hydrology in indian himalayan region: a review. Wiley Interdisciplinary Reviews: Climate Change, 7(3):393–410, 2016.

A. Staal, S. C. Dekker, C. Xu, and E. H. van Nes. Bistability, spatial interaction, and the distribution of tropical forests and savannas. Ecosystems, 19(6):1080–1091, 2016.

A. Staal, I. Fetzer, L. Wang-Erlandsson, J. H. Bosmans, S. C. Dekker, E. H. van Nes, J. Rockström, and O. A. Tuinenburg. Hysteresis of tropical forests in the 21st century. Nature communications, 11(1):1–8, 2020.

A. C. Staver, S. Archibald, and S. A. Levin. The global extent and determinants of savanna and forest as alternative biome states. science, 334(6053):230–232, 2011.

J. T. Tanzil, B. E. Brown, R. P. Dunne, J. N. Lee, J. A. Kaandorp, and P. A. Todd. Regional decline in growth rates of massive p orites corals in s outheast a sia. Global change biology, 19(10): 3011–3023, 2013.

A. T. Vanak, G. Shannon, M. Thaker, B. Page, R. Grant, and R. Slotow. Biocomplexity in large tree mortality: interactions between elephant, fire and landscape in an african savanna. Ecography, 35(4):315–321, 2012.

O. Venter, E. W. Sanderson, A. Magrach, J. R. Allan, J. Beher, K. R. Jones, H. P. Possingham, W. F. Laurance, P. Wood, B. M. Fekete, et al. Global terrestrial human footprint maps for 1993 and 2009. Scientific data, 3(1):1–10, 2016.

J. Xu, R. E. Grumbine, A. Shrestha, M. Eriksson, X. Yang, Y. Wang, and A. Wilkes. The melting himalayas: cascading effects of climate change on water, biodiversity, and livelihoods. Conservation Biology, 23(3):520–530, 2009.

X. Xu, D. Medvigy, A. T. Trugman, K. Guan, S. P. Good, and I. Rodriguez-Iturbe. Tree cover shows strong sensitivity to precipitation variability across the global tropics. Global ecology and biogeography, 27(4):450–460, 2018.

P. Yadava. Savannas of north-east india. Journal of biogeography,pages 385–394, 1990.

L. G. Ziccardi, M. Dos Reis, P. M. L. d. A. Graça, N. B. Gonçalves, A. Pontes-Lopes, L. E. Aragão, R. P. de Oliveira, L. Clark, and P. M. Fearnside. Forest fires facilitate growth of herbaceous bamboos in central amazonia. Biotropica, 2021.

