## Supplementary File for "Characterising the vegetation-rainfall relationship in the Northeast Himalaya, India"

#### Supplementary 1: Human Footprint index and GlobCover map

To remove the confounding factors such as agriculture fields and built-up area, we masked the global land cover map (2009) with the human footprint index map (HFI) (Bontemps et al. 2011). We selected agricultural fields, tea gardens, built-up area, and forest to examine the associated HFI values and the corresponding landcover type. A total of 220 points were manually verified. The HFI values for northeast India ranged from 0 to 43. Global land cover type map showed that natural forest area corresponds to HFI value of 0–8, whereas the agriculture field, tea garden area and built-up areas corresponded to HFI >8. Therefore, we selected '8' as a cut-off (threshold

Current address: Centre for Ecological Sciences, Indian Institute of Science, Bengaluru

limit) to remove the pixels with high human influence. Pixels with values $\leq 8$ were retained for further analysis, while those with $> 8$ were removed. Pixels associated with high human footprint are shown in Fig. 1.

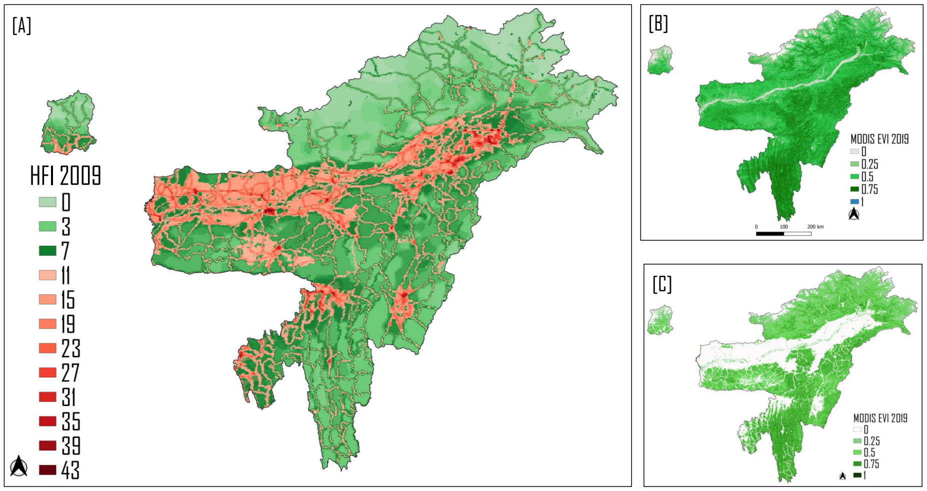


**Figure 1:** [A]. Human footprint index for northeast India. HFI values range from 0-43; pixels with a value of 43 show highest human footprint [B] spatial distribution of Enhanced Vegetation Index (EVI) without removing the human footprint area, and [C] after removing the human footprint filtering out pixel >8

#### Supplementary 2: Influence of Human footprint index in state diagram

We constructed a state diagram without removing the pixels with high human footprint to examine if they influenced our results. We plotted the Enhanced Vegetation Index (EVI) (without removing human footprint) as a function of Mean Annual Precipitation (MAP). The plot was consistent with our previous result (when pixels >8 were removed); EVI with the increased with MAP till 2000 mm (Fig. 2).

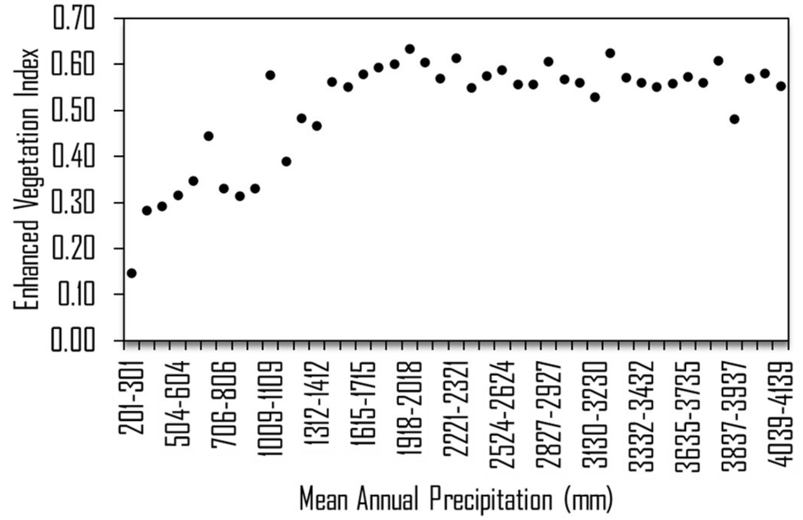


**Figure 2:** Ecosystem state diagram i.e., EVI (with human footprint) as a function of MAP (at 100 mm bin) show mode of EVI increases gradually along with the MAP and horizontally asymptote at >2000 MAP

#### Supplementary 3: Construction of state diagram using mean EVI values

We constructed a state diagram using mean values of EVI (using the function mean in R) as a function of MAP. The resulting state diagram was similar to our previous result, with EVI increasing with MAP till 2000 mm (Fig. 3).

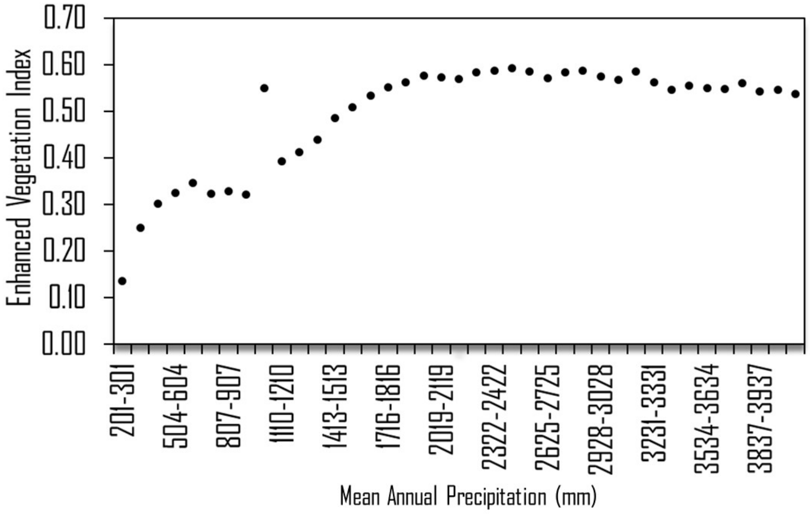


**Figure 3:** Ecosystem state diagram i.e., mean of EVI as a function of MAP show EVI gradually increase along with the MAP and asymptote at >2000 mm MAP.

#### Supplementary 4: Relationship between elevation, mean annual precipitation and surface temperature

A scatter plot was used to analyse the relationship between temperature and elevation, as well as precipitation and elevation. We obtained data for elevation (SRTM DEM, at 30 m resolution), median temperature (MODIS Terra Land Surface Temperature and Emissivity 8-Day Global product, at 1km resolution) and mean annual precipitation (CHIRPS data, at 1km resolution) (Wan et al. 2006; Funk et al. 2015). Three transects, two spanning regions in Arunachal Pradesh (transects 1 and 2) and 1 in Nagaland (transect 3), were drawn across the NEI region (Fig. 4). We collected data of elevation, MAP and temperature for points within the transects for the period January 2018 - January 2019. The results of the analysis showed that median surface temperature and mean annual precipitation decreased with increase in elevation gradient. All analyses were performed in Google Earth Engine.

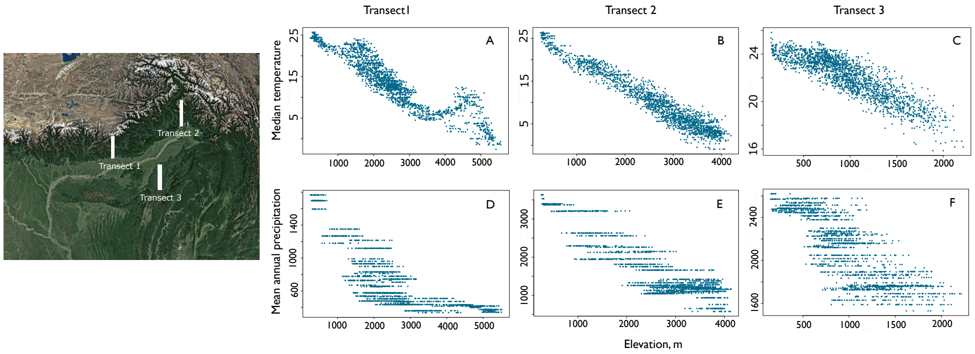


**Figure 4:** Over the elevation, a scatter plot of median temperature (°C) and mean annual precipitation (mm) for the three transect (transect 1, 2 and 3); with increasing elevation, the median temperature (A, B and C) and mean annual precipitation decreases (D, E and F)

### References


S. Bontemps, P. Defourny, E. Van Bogaert, O. Arino, V. Kalogirou, and J. R. Perez. Globcover 2009-products description and validation report. *URL: http://ionia1. esrin. esa. int/docs/GLOBCOVER2009_Validation_Report_2*, 2, 2011.

C. Funk, P. Peterson, M. Landsfeld, D. Pedreros, J. Verdin, S. Shukla, G. Husak, J. Rowland, L. Harrison, A. Hoell, et al. The climate hazards infrared precipitation with stations—a new environmental record for monitoring extremes. *Scientific data*, 2(1):1–21, 2015.

Z. Wan et al. Modis land surface temperature products users' guide. *Institute for Computational Earth System Science, University of California: Santa Barbara, CA, USA*, 805, 2006.
